# Advancing the quantification of land-use intensity in forests: the ForMIX index combining tree species composition, tree removal, deadwood availability, and stand maturity

**DOI:** 10.1101/2025.03.10.642318

**Authors:** Michael Staab, Nico Blüthgen, Katja Wehner, Peter Schall, Christian Ammer

## Abstract

Many forests have a long history of human land use, which relates to species communities and ecosystem processes, making robust and quantitative measures of land-use intensity in forests desirable. We here introduce the ForMIX (Forest Management IndeX), a compound index combining altered tree species composition, tree removal, deadwood availability and stand maturity, which are each calculated as the deviation from expectations in a natural old-growth forest reference. By relating to resources and niches directly affected by forest land use, the compound index and its components allow for mechanistic inference on the consequences of land use in forests. Using basic forest inventory data from 150 sites distributed over three regions of Germany, we demonstrate the properties of ForMIX, which differentiates well among forest types and silvicultural systems and is robust to decisions regarding reference values and components. Reference values used in ForMIX are dynamic, may shift with ongoing climate change and may require refinement for different geographic regions. ForMIX advances the quantification of land-use intensity in forests by being biologically meaningful, being usable and comparable across forest types, being derivable from standard forest inventory data, and by being easy to apply, understand and interpret.

## Introduction

Forests are ecologically and economically important ecosystems (Brockerhoff et al. 2017). Covering around one third of global land area, forests provide habitat for a large proportion of global biodiversity and are pivotal for the global carbon cycle (FAO and UNEP 2020; Pan et al. 2024). At the same time, forests support the livelihoods of a sizeable share of mankind. This is why most forests, particularly in temperate climates, have a long history of human land use, which alters forest ecosystems and influences species communities and ecological processes (Hansen et al. 2013; Penone et al. 2019). Since human interventions range from complete clear cutting of trees in short rotation periods to rather extensive forms of forest management, such as single tree selection within the concept of continuous cover forestry (Bettinger et al. 2016; Pommerening 2024), robust measures to quantify land use in forests are desirable. Such measures then allow to study influences of forest management on biodiversity, ecological processes and multifunctionality, and may also guide policy, for example when deciding which forests should be set aside in conservation programs.

Several approaches to measure and quantify land-use intensity are applied to forests (reviewed in Bellamy et al. 2024; Schall and Ammer 2013). For example, the presence or abundance of specific indicators (Caro 2010), such as beetles associated with old growth forest (primeval forest relict species, Eckelt et al. 2017), can inform on past land use. Other approaches rely on the identity and quantity of specific tree-related microhabitats (Larrieu et al. 2018), forest naturalness or disturbance (Senf and Seidl 2021). A range of compound indices (Greco et al. 2019) that combine different independent raw measurements into a single value, are also available to describe land use in forests (e.g. Kahl and Bauhus 2014; Schall and Ammer 2013; Storch et al. 2018). Any measure of land-use intensity in forests should have several desirable properties if it is to be applied in ecological analyses beyond a specific local context: the measure should be biologically meaningful and ideally relate to resources and niches on which forest organisms depend and which are altered by land use; it should be easy to understand and interpret; it should be comparable among different forest types and abiotic conditions; it should quantitatively distinguish different management scenarios; it should be readily computable from basic forest inventory data to facilitate wide dissemination and application.

Two examples for such measures are the Forest Management index (ForMI, Kahl and Bauhus 2014) and the Silvicultural Management Intensity indicator (SMI, Schall and Ammer 2013), which were both conceived for temperate forests in Central Europe. ForMI is calculated as the sum of three components: the volume proportion of non-natural tree species, the ratio of harvested tree volume (based on remaining stumps) in relation to the sum of standing, harvested and deadwood volume, and the proportion of dead wood volume with saw cuts in relation to the total deadwood volume (Kahl and Bauhus 2014). SMI, in turn, consists of two components in comparison to reference data from yield tables: the basal area in a forest stand, and the risk of stand loss due to management, which both vary among tree species and depend on actual stand age and hence the applied management regime (Schall and Ammer 2013). Indices generally trade off complexity, applicability and interpretation (among other considerations), and no index is without constraints (Greco et al. 2019). Both, ForMI and SMI differentiate well between managed and unmanaged stands, but have several perceived shortcomings when used in ecological analyses. From a resource perspective, the origin of deadwood (sawing or natural) in ForMI should not matter for deadwood-associated consumer species. Also, the necessary data on deadwood origin are not readily recorded in forest inventories. The ratio of harvested trees in ForMI is based on volume, requiring tree height measurements in addition to tree diameter for allometries. This component scores stumps of any decay stage (sensu Müller-Using and Bartsch 2009; Siitonen 2001), meaning that trees cut several decades ago are considered, even though the disturbance and resource removal associated to tree harvest is likely to have compensated earlier due to regrowth and subsequent canopy closure. SMI, in turn, is rather abstract and the compound index or its components are difficult to interpret ecologically. Moreover, the density component in SMI is based on yield tables derived in pure stands which do only partly apply to mixed stands.

To further advance the quantification of land-use intensity in forests, we here propose the ForMIX (Forest Management IndeX) index that blends the components (1) altered tree species composition, (2) tree removal, (3) deadwood availability, and (4) stand maturity. All four components are calculated as deviations from a reference forest ecosystem in the absence of land use. By default, the components range between 0 (no use) and 1 (theoretical maximum use) and can simply be summed to the new compound index. As the components are not strongly correlated, they capture independent aspects of land use and forest management. The new index can be easily computed from standard inventory data, is comparable across forests types, and easy to interpret. If adjusted to regional reference systems, it can readily be applied to any forest.

## Materials and Methods

### Rationale of the new ForMIX index

When first conceiving the ForMIX, the aim was to find a measure that allows a biological interpretation of land-use intensity in forest by ideally building on resources and niches that are important for forest organisms and at the same time affected by forest management. Further considerations were easy understanding, comparability, and straightforward calculation with only basic forest inventory data needed. The ForMIX is a compound index consisting of the four additive components *tree species composition*, *tree removal*, *deadwood availability*, and *stand maturity* (Table 1) that each represent different dimensions of how land-use intensity relates to resources and niches, partly building on the rationale originally applied by Kahl and Bauhus (2014). All components are quantitative and consequently assess the local deviation from a natural old-growth forest in terminal stage without any use. Components range between 0 (no use) and 1 (theoretical maximum). ForMIX is simply the sum of the four components, with a low value thus indicating a forest little influenced by human activity, while a high value indicates an intensively used forest.

**Table 1.**
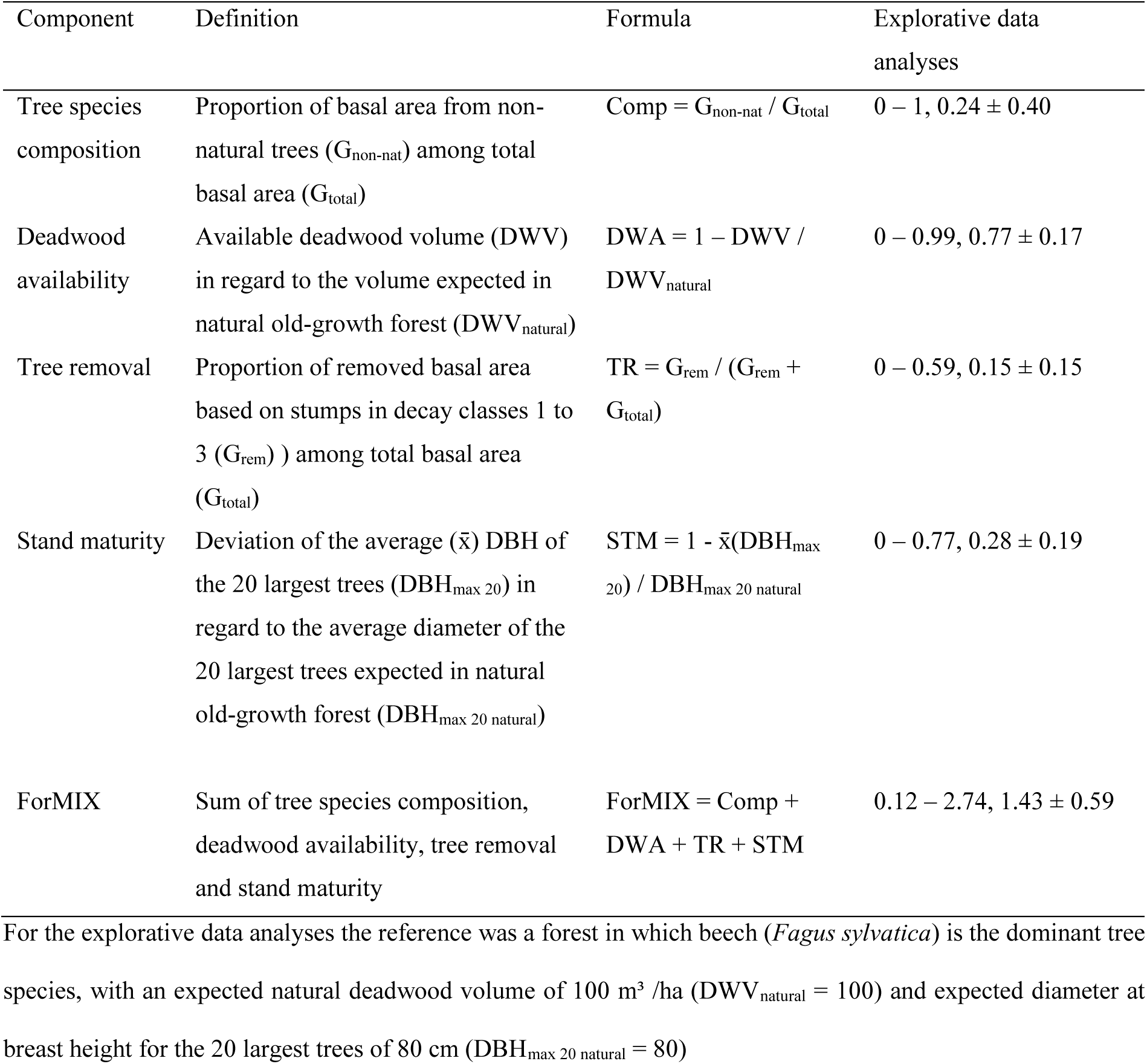
Overview, definitions and formulae of the ForMIX index. Explorative data analyses were conducted for 150 forest stands from three regions of Germany. Values give the range (min – max) and the mean ± SD

It should be noted that each component can also be used individually in a meaningful way to explore mechanisms behind variation in effects of forest use. Three components represent changes in biotic resources and environmental conditions from the viewpoint of organisms (*tree species composition*, *deadwood availability*, *stand maturity)* whereas *tree removal* represents both a classic disturbance (albeit infrequent) and altered conditions such as light regime. The four components are assumed to have complementary effects on communities that vary across species.

*Tree species composition* (Comp) is the proportion of basal area from trees of species (G_non-nat_) among total basal area (G_total_) that do not belong to the regional set of tree species occurring under present natural conditions, calculated as Comp = G_non-nat_ / G_total_. In the majority of Central European forests, the species identity of trees is a direct outcome of past forest management (Stark et al. 2021), and, hence, the component describes how much tree species composition has been altered. Which tree species could potentially occur naturally at a given site depends on climate, elevation and soil, among other site characteristics. Suitable site-specific reference composition can, for example, be derived from the European Forest Types (Barbati et al. 2007, Barbati et al. 2014) or national forest inventories (e.g. for Germany: https://bwi.info/). However, trees with favorable attributes have often been propagated in silviculture. This has led to situations where forests are partly dominated by tree species that would not occur naturally at these sites. Examples in Central Europe are forests in lowlands with Norway spruce (*Picea abies*), European larch (*Larix decidua*), Scots pine (*Pinus sylvestris*), or Douglas fir (*Pseudotsuga menziesii*). While Douglas fir originates from North America, and is hence clearly not native, Norway spruce, European larch and Scots pine are native to Central Europe, but would, without propagation by forestry, be restricted to high elevations (spruce, larch) or specific climates and soils (pine). Thus, these important timber species are considered not natural for most of Central Europe, where the dominating potential natural vegetation outside mountains, wetlands and very dry soils is a broadleaved forest with a high share of European beech (*Fagus sylvatica*) (Barbati et al. 2007; Suck et al. 2014; Tüxen 1956). Non-natural trees directly link to resources and relate to many ecological processes. When their density is high, non-natural trees can influence the community composition of consumer species across trophic levels (Penone et al. 2019; Purahong et al. 2018; Staab et al. 2021; Wildermuth et al. 2024).

*Deadwood* availability (DWA) is the deviation of the deadwood volume (DWV) in regard to the volume expected in natural old-growth forests (DWV_natural_), calculated as DWA = 1 – DWV / DWV_natural_. Deadwood is a key resource in forests, on which very many forest species depend for food and habitat (Stokland et al 2012). Furthermore, deadwood retains moisture and is central for the carbon cycle in forests. One net outcome of forestry is lower deadwood availability than in unmanaged old-growth forest (Müller and Bütler 2010), because trees are being felled before they develop larger amounts of deadwood due to natural mortality, or because most deadwood is actively removed before entering decay, for example in sanitary cuttings. In Central Europe, a natural beech forest would in mean accumulate approximately 100 m³/ha deadwood (Commarmot et al. 2013; Nagel et al. 2023), which is proposed as reference (see considerations on the choice of reference values below). If a forest stand has more deadwood per area than the reference, the calculation results in a negative value, which is set to 0 (optimal value).

*Tree removal* (TR) is the proportion of basal area of all stumps of trees that had been removed (G_rem_) among total basal area (G_total_) including living trees, calculated as TR = G_rem_ / (G_rem_ + G_total_). Removing trees from a forest, either due to final harvest or due to other interventions such as thinning, is the main difference between forest management and natural disturbances (Meyer and Ammer 2022). Once a tree has been taken from a stand, the resource and habitat structures provided by this tree are lost, the local soil and vegetation may be disturbed, and the canopy opens, which changes the microclimate (Geiger 1965). Over time, resources regrow, disturbance diminishes, and the canopy closes, which largely compensates for the effects of tree removal. For the calculation of the tree removal component, we propose to include stumps from decay classes 1 to 3 (sensu Müller-Using and Bartsch 2009; Siitonen 2001), which accounts for trees removed in the last 10 to 15 years, the period of time after which the direct consequences of removing trees will largely have disappeared, at least in the selection cutting forestry conducted in Central Europe. If temporally resolved and stand-specific tree removal data recorded at the same spatial scale as the data for the other ForMIX components are available, these could be used instead of stumps to calculated tree removal. We note that TR might slightly overestimate the actual removal if basal area of living trees is calculated based on DBH (diameter at breast height) data, as the diameter at the level of a stump is slightly larger than at breast height.

*Stand maturity* (STM) is the deviation of the average (x̄) DBH of the 20 largest trees per ha (DBH_max 20_) in regard to the average diameter of the 20 largest trees per ha expected in natural old-growth forest (DBH_max 20 natural_), calculated as STM = 1 - x̄(DBH_max 20_) / DBH_max 20 natural_. For most tree species large trees are commercially more valuable and are usually harvested before reaching their maximum natural size (Faustmann 1849). Hence, a lack of large and old trees is a characteristic of land use in forests. Large trees are important for many forest-associated species, because they provide resources and (micro-)habitats not available from smaller trees (Müller et al. 2014), which is why large trees and older forests often sustain higher species diversity (e.g. Lutz et al. 2018; Moning and Müller 2009). Furthermore, large trees buffer forest microclimate (Frey et al. 2016; Lindenmayer et al. 2022). As reference for the calculation of the stand maturity component, the diameter of the 20 largest trees is proposed, as this number can easily be subdivided, without resulting in decimal numbers, when inventory area is smaller than 1 ha. The 20 largest trees in natural Central European broadleaved forests forest would at most sites have a mean diameter of approximately 80 cm (e.g. Hobi et al. 2015; Vandekerkhove et al. 2018), which is here used as reference for DBH_max 20 natural_. In other forest types this value could obviously differ. For example, for hemiboreal and subalpine forests, 40 cm would be a more appropriate reference. If the mean diameter of the 20 largest trees per ha is larger than the reference, the calculation results in a negative value, which set to 0 (optimal value), analog to deadwood availability.

The four components are simply summed to receive the ForMIX (ForMIX = Comp + DWA + TR + STM), with higher values indicating higher land-use intensity. One issue to consider for any aggregated compound index is standardization of the components (Greco et al. 2019). In case of the ForMIX, for example, a tree removal of 1 is only possible in completely clear-cut stand with no living trees left. Likewise, a stand maturity of 1 is only theoretically possible when all trees are seedlings or saplings. If it is desirable to ensure an equal contribution of all components in the compound index, each component may be standardized between 0 (minimum in data) and 1 (maximum in data) before the compound index is calculated (see also Blüthgen et al. 2012). However, be aware that doing so may overemphasize a single component when the range between minimum and maximum in this component is small.

If geographically and climatically appropriate reference values are used in calculations of components, the consequent referral to the expectations in natural forests without land use allows to apply the ForMIX across a wide range of forest types. This can facilitate comparative analyses beyond local context, for example in forest inventories at the regional or even national level. To test the ForMIX (see below), we used data from sites that would naturally be dominated by beech and we here apply reference values for altered tree species composition, deadwood availability and stand maturity corresponding to beech forest, as this would be the prevailing forest type for the majority of Central Europe. Nonetheless, there would also under natural conditions be many forest types other than beech in Central Europe, such as mountain forests dominated by conifers (with naturally high proportions of spruce and larch) or dry forests with a high proportion of oak and a natural contribution of pine (Suck et al. 2014; Tüxen 1956). By using forest type-specific references (e.g. derived from national forest inventories) in the simple formulas of the respective ForMIX components, the index can easily be transferred to basically any forest. For Europe, the European Forest Types (Barbati et al. 2007; Barbati et al. 2014) could be a suitable unifying framework. If similar data for expectations under conditions without land use exist, ForMIX can of course also be applied beyond Central Europe.

### Forest inventory data

To parameterize and test the ForMIX, we used forest inventory data from a total of 150 sites (size 1 ha, 100 m x 100 m) distributed across the three German regions Schwäbische Alb, Hainich-Dün and Schorfheide-Chorin (Fig. S1, Supplementary material) that are part of the Biodiversity-Exploratories framework (www.biodiversity-exploratories.de, Fischer et al. 2010). The regions cover a range of soil types, topography, precipitation and mean annual temperature, making them representative for large parts of Central Europe. Sites in each region were originally chosen out of a large pool of potential candidate sites by a stratified procedure and include a broad range of forest types, from intensively managed planted conifer stands (Norway spruce: *Picea abies*, 16 sites; Scots pine: *Pinus sylvestris*, 20 sites) to managed stands of European beech (*Fagus sylvatica*, 82 sites) and oak (*Quercus* sp., 7 sites). Unmanaged natural broad-leaved forests are also part of the study design (25 sites). The tree layer in the potential natural vegetation on all sites would under the criteria of the German national forest inventory be dominated by beech. While being native to Central Europe, under natural conditions the planted conifers would on the sites be absent (pine) or only occur at low densities in the higher elevations of Schwäbische Alb (spruce).

On each 1 ha site, a full forest inventory was conducted between 2015 and 2018 (completed before the drought of summer 2018). Every living tree with a diameter at breast height (DBH) larger 7 cm was identified to species and its DBH recorded. Deadwood items, including stumps, larger 25 cm diameter were measured (diameter and length) at the entire site, while smaller deadwood (7-25 cm diameter) was measured across the site diagonals on two 2 m-wide line transects that were subsequently extrapolated to 1 ha. The decay stage (Müller-Using and Bartsch 2009; Siitonen 2001) of each deadwood item was classified on an ordinal scale between 1 (fresh deadwood) and 5 (heavily decayed). More details on site selection, forest characteristics, inventory procedures and local forest management practices are available in Fischer et al. (2010), Schall et al. (2018) and Staab et al. (2023).

### Explorative data analyses

Using these basic inventory data, the ForMIX index was calculated as the sum of the four components altered tree species composition, deadwood availability, tree removal, and stand maturity applying the formulas described above. According to the German national forest inventory, beech (*Fagus sylvatica*) would be the main tree species at all sites. Hence, the reference for the index calculation in the explorative data analyses was a beech forest, with an expected deadwood volume of 100 m³/ha under natural conditions and an expected 20 trees with DBH larger 80 cm per ha. Regarding tree species composition, beech would in each of the three regions be accompanied by a set of region-specific subordinate tree species and pioneer tree species. All trees that are neither from main, nor subordinate nor pioneer tree species are considered non-natural. If conifer species are subordinate but account for more than 1/3 of basal area (i.e. occurring at densities that are not subordinate anymore), the species are also considered non-natural, because they have been propagated by forestry in the past and because they are phylogenetically distant to the otherwise naturally occurring broadleaved tree species. In our data, this is the case for spruce monocultures in Schwäbische Alb, where Norway spruce is considered a subordinate tree species due to the high elevation of the sites.

All analyses were conducted in R 4.3.1 (www.r-project.org). To assess relationships or potential non-independence among the four components, Spearman correlation coefficients were calculated. To test if the ForMIX differentiates among the forest types occurring in the inventory data (managed spruce, pine, beech, and oak forest as well as unmanaged forest) linear mixed-effects models were calculated with the R-package lme4 (Bates et al. 2015). Response variables were the ForMIX or its separate components with the categorical forest type being the only fixed effect. Region was treated as random intercept to account for regional differences in forest characteristics and forestry practices. Significance (using Kenward-Roger approximated degrees of freedom) among the categorical forest types was evaluated with the R-package emmeans (Lenth 2020) by applying the Tukey method to account for possible type I errors. We also calculated a ForMIX in which all components were standardized between 0 (minimum in raw component) and 1 (maximum in raw component). The Spearman correlation between raw and standardized ForMIX was assessed. To provide an example with alternative reference values (other than beech forest) in the components, ForMIX was also computed under the fictive assumption that spruce, larch and pine are native at all sites independent of their contribution to total basal area and that the expected amount of deadwood is 60 m³ / ha. The Spearman correlation between the regular ForMIX, which uses beech forest as reference, and the fictive alternative ForMIX was calculated. Finally, ForMIX was compared (Spearman correlation) with the Forest Management Intensity index (ForMI, Kahl and Bauhus 2014) and the Silvicultural Management Intensity indicator (SMI, Schall and Ammer 2013), two alternative measures of land-use intensity in forests.

## Results

The ForMIX index applied to forest inventory data from 150 sites across three regions of Germany took values from 0.12 to 2.74 (mean 1.43 ± 0.59 SD, Table 1). The compound index, calculated as the sum of the components altered tree species composition (range 0-1, mean 0.24 ± 0.40 SD), deadwood availability (0 – 0.99, 0.77 ± 0.17), tree removal (0 – 0.59, 0.15 ± 0.15), and stand maturity (0 – 0.77, 0.28 ± 0.19), was approximately normally distributed, and leveled the distribution of its single components (Fig. 1). Tree species composition, deadwood availability and tree removal were not correlated among each other, while stand maturity was correlated to the three other components (Fig. 2). Most importantly, ForMIX differentiated well among forest types (Fig. 3), with managed pine and spruce forests achieving highest ForMIX, managed beech and oak intermediate ForMIX, and unmanaged forests lowest ForMIX (statistical details in Table S1, Supplementary material). This clear distinction between management types was not recovered with any component alone. For example, the composition of spruce and pine forests was obviously dominated by non-natural trees, while deadwood availability was largely similar across forest types (Fig. 3). ForMIX was indifferent to standardization (Fig. S2, Supplementary material), with the index based on standardized components being highly correlated to the regular index based on raw components (Spearman’s rho=0.993, p<0.001). ForMIX was also robust to alterations in the reference values of its components. A fictive alternative treating spruce, larch and pine as natural on all sites and using a lower reference for deadwood availability (Fig. S3, Supplementary material) was highly correlated (Spearman’s rho=0.824, p<0.001) to regular ForMIX using beech forest as reference. Furthermore, ForMIX was highly correlated to both ForMI and SMI (Fig. S4, Supplementary material).

**Fig. 1.**
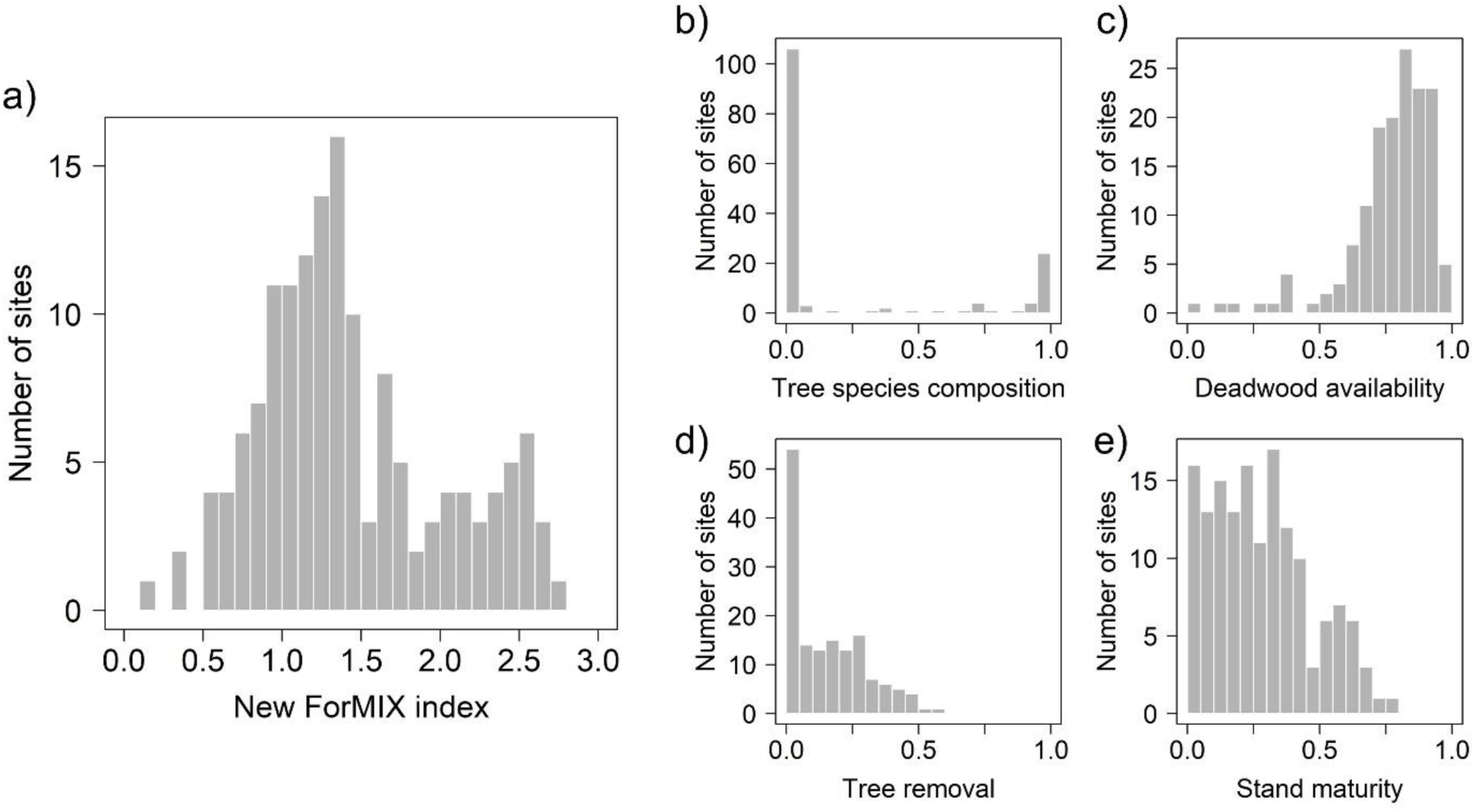
Histograms illustrating the distribution of a) the new ForMIX index and the components b) tree species composition, c) deadwood availability, d) tree removal, and e) stand maturity from which the index is calculated. Data are based on forest inventories conducted on a total of 150 forest sites (size 1 ha) distributed over three regions of Germany (Fig S1, Supplementary material)

**Fig. 2.**
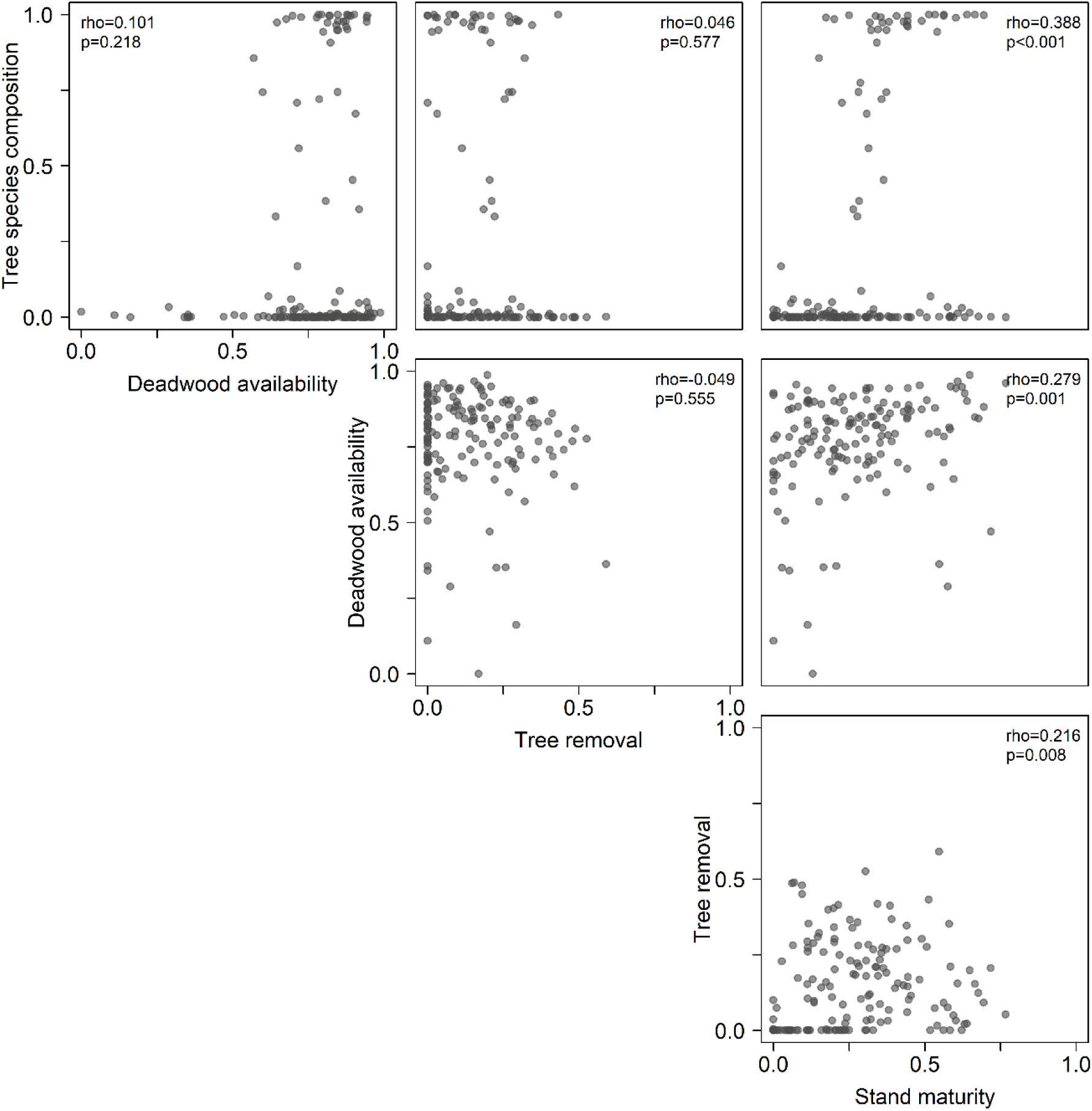
Scatterplots illustrating relationships among the components (tree species composition, deadwood availability, tree removal, stand maturity) of the ForMIX index. Insets give Spearman correlations (rho) and corresponding p-values. Only stand maturity is related to other components. Data are based on forest inventories conducted on a total of 150 forest sites (size 1 ha) distributed over three regions of Germany (Fig S1, Supplementary material)

**Fig. 3.**
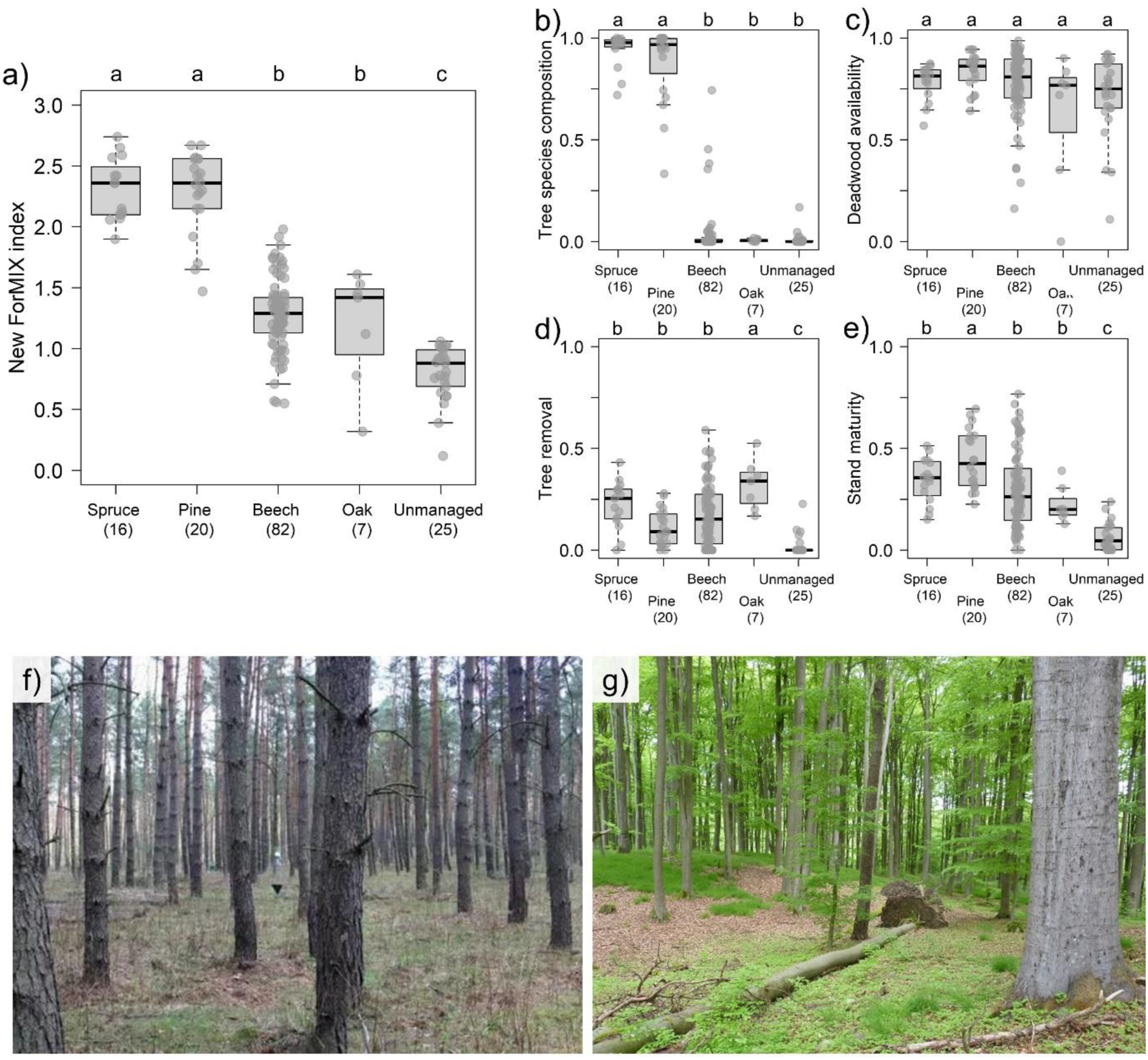
The new ForMIX index a) differentiates among forest types (managed spruce, pine, beech, oak forest as well as unmanaged forest). b-e) show the components on which the index is based. Letters above boxplots indicate grouping of estimates (based on the Tukey method), with alphabetically decreasing group means (see Table S1 for exact pairwise contrasts). f) is an example for an intensively managed pine forest with high ForMIX = 2.56 and g) for an unmanaged forest with low ForMIX = 0.69. Data are based on forest inventories conducted on a total of 150 forest sites (size 1 ha) distributed over three regions of Germany (Fig. S1, Supplementary material). Pictures in f) and g) by Michael Staab

## Discussion

The new ForMIX index has several favorable properties facilitating the analyses of land-use intensity in forests. ForMIX not only distinguishes among forest types with different sylvicultural systems, as do other indices (e.g. Kahl and Bauhus 2014; Schall and Ammer 2013). Because its components are directly derived from deviations in resources and environmental conditions affected by land-use intensity in forests, the index is also biologically interpretable and allows for mechanistic inference. This also applies to each of its individual components (tree species composition, deadwood availability, tree removal, stand maturity). Importantly, by being computable from basic forest inventory data without requiring elaborated mathematical or statistical knowledge, by being easy to understand and being comparable across forest types, ForMIX can be applied in forest ecology beyond specific local context. While described and applied for beech forest in our study, i.e. the potential natural vegetation in large parts of Central Europe, the rationale behind the ForMIX remains flexible. By adapting the references for non-natural tree species (i.e. species that under present natural conditions do not belong to the regional set of tree species), deadwood volume and diameter of large trees, for example using the framework of the European Forest Types (Barbati et al. 2007; Barbati et al. 2014), the ForMIX index can in principle be used to quantify land-use intensity in any forest. For simplicity, the components deadwood availability and stand maturity use an area of 1 ha as reference, which can, however, by multiplication or division be easily upscaled or downscaled to the size of the inventoried area (e.g. 25 m³ /ha deadwood and 5 trees larger 80 cm DBH when the inventory area is 0.25 ha).

The new index has several further advantages, even though other land-use intensity indices in forests (ForMI, Kahl and Bauhus 2014; SMI, Schall and Ammer 2013) were correlated with ForMIX. Due to the consequent referral to regional references in forests without land use, yield tables or other abstract approximations are not necessary. By deriving the components from basal area instead of volume, no allometries are applied and measurements of tree height, which are not always available in basic inventories, are not required. However, with the increasing availability of terrestrial laser scanning for forest inventories (Ehbrecht et al. 2017; Liang et al. 2016), it may be that height and other parameters will soon become more readily available. ForMIX could then easily use volume instead of basal area. While compound indices are favored by many users (Greco et al. 2019), and the ForMIX allows direct inference regarding land use and its intensity, the single components can in many situations be biologically informative, as the components capture independent aspects of resources and niches and, hence, connect to mechanisms. Therefore, using all four ForMIX components jointly as covariates in regression models is possible, as the components are largely uncorrelated. Exceptions are the correlations with stand maturity, which are expected, as non-natural trees propagated by forestry are harvested earlier, older forests have necessarily not been cut in the past, and as stand age is central to the ecology of a stand (Hilmers et al. 2018), with older forests having, for example, more deadwood. All correlation coefficients among components are below thresholds that indicate problems of collinearity (Dormann et al. 2013), but users are encouraged to check for potential influences on variances (e.g. with variance inflation factors) if all four components are used as covariates in the same statistical model.

As for any other index, there are scenarios where the application of ForMIX will not be possible, for example after a major natural disturbance such as large-scale windthrow, fire or an insect calamity, which will change the components without being an outcome of land use. Also, ForMIX may miss certain forestry actions, with one example being tree removal if stumps are completely removed after harvest, a practice that is no longer regularly conducted in Central Europe. Many forests worldwide are now being invaded by non-native tree species (Delavaux et al. 2023). If these non-native trees were not planted but invaded a forest by natural dispersal, they would still be counted in the ForMIX component tree species composition, although not directly being directly an outcome of land use.

Except for species that clearly originate from other continents, there is ongoing discussion about the naturalness of tree species used in forestry (Stark et al. 2021). In some regions, including the lowlands of Central Europe, currently important timber species would not occur naturally at many sites (even though originating from the region), but have been planted in silviculture for centuries (Müller et al. 2024). While the respective species (especially Norway spruce, *Picea abies*, and Scots pine, *Pinus sylvestris*) are usually treated as not natural, they support an associated flora and fauna (Brändle and Brandl 2002; Leidinger et al. 2019). If a user of ForMIX is concerned that in forests containing many non-natural trees, the tree species composition component could have a too large contribution to the compound index, this component could potentially be weighted down by multiplication with a constant >0 and <1, or even be excluded from analysis. We basically explored this option in the explorative data analyses, by treating spruce, larch and pine as natural in all sites, but the resulting fictive alternative ForMIX was still highly related to the regular ForMIX. Similarly, standardization of all components resulted in a version of ForMIX that was strongly correlated with the regular index using raw components. Taken together, these alternative calculations provide further evidence for the robustness of the rationale behind ForMIX that is unlikely to be fundamentally biased by single decisions regarding components and reference values.

One strength of the ForMIX is to consequently assess forest land-use intensity as the deviation from a natural reference system. Notwithstanding, with ongoing climate change it will in the future become inevitable to refine reference values for all components except tree removal. Tree species are expected to change their distribution and abundance in complex and not yet predictable ways (Laughlin and McGill 2024; Wessely et al. 2024). For example, it is unclear if forests dominated by European beech (*Fagus sylvatica*) continue to be the potential natural vegetation in Central Europe (Martinez del Castillo et al. 2022; Gessler et al. 2024). Likely, by the end of the century, forests in Europe and beyond will also under natural conditions have changed, requiring new definitions of the natural forest types (sensu Barbati et al. 2007; Suck et al. 2014). Consequently, a future ForMIX might treat different tree species as not natural and might be based on altered reference values for deadwood volume and diameter of large trees. Nevertheless, this future ForMIX would still be a biologically meaningful measure of land use in forests that is comparable among forest types, can directly be calculated from standard forest inventory data, and is easy to understand and interpret.

## Availability of data and material

This work is based on data elaborated by projects of the Biodiversity Exploratories program (DFG Priority Program 1374). The dataset is publicly available in the Biodiversity Exploratories Information System (http://doi.org/10.17616/R32P9Q) under the accession number 31855.

## Authors’ contributions

M.S. conceived the idea for the index, which was refined with support from all coauthors; P.S. and C.A. provided forest inventory data; M.S. analyzed the data and led the writing. All authors revised drafts and gave final approval for publication.

## Acknowledgements

We thank the managers of the three Exploratories, Martin Gorke, Katrin Lorenzen, Kirsten Reichel-Jung, Miriam Teuscher, Juliane Vogt, and all former managers for their work in maintaining the plot and project infrastructure; Christiane Fischer and Jule Mangels for giving support through the central office, Michael Owonibi and Andreas Ostrowski for managing the central data base, and Markus Fischer, Eduard Linsenmair, Dominik Hessenmöller, Daniel Prati, Ingo Schöning, François Buscot, Ernst-Detlef Schulze, Wolfgang W. Weisser and the late Elisabeth Kalko for their role in setting up the Biodiversity Exploratories project. Jörg Memmert provided data for maps. We thank Franz Kroiher for valuable discussion on the German National Forest Inventory, and the administration of the Hainich national park, the UNESCO Biosphere Reserve Swabian Alb and the UNESCO Biosphere Reserve Schorfheide-Chorin as well as all land owners for the excellent collaboration. Field work permits were issued by the responsible state environmental offices of Baden-Württemberg, Thüringen, and Brandenburg.

