## Supplemental Information for "Advancing the quantification of land-use intensity in forests: the ForMIX index combining tree species composition, tree removal, deadwood availability, and stand maturity"

**Table S1** Pairwise contrasts of ForMIX and its components between forest types (managed spruce, pine, beech, oak forest as well as unmanaged forest) for 150 sites in three regions of Germany (Fig. S1). P-values are derived from the Tukey-method and based on Kenward-Roger approximated degrees of freedom. Significant contrasts (at $p < 0.05$) are shown in bold

| Contrast | Estimate ± SE | *t*-ratio | p-value |
|---|---|---|---|
| *ForMIX* | | | |
| Spruce – Pine | -0.104 ± 0.128 | -0.807 | 0.928 |
| **Spruce – Beech** | **0.977 ± 0.086** | **11.316** | **<0.001** |
| **Spruce – Oak** | **0.997 ± 0.156** | **6.379** | **<0.001** |
| **Spruce – unmanaged** | **1.428 ± 0.103** | **13.819** | **<0.001** |
| **Pine – Beech** | **1.080 ± 0.096** | **11.301** | **<0.001** |
| **Pine – Oak** | **1.101 ± 0.129** | **8.556** | **<0.001** |
| **Pine – unmanaged** | **1.532 ± 0.104** | **14.725** | **<0.001** |
| Beech – Oak | 0.021 ± 0.131 | 0.158 | 1.000 |
| **Beech – unmanaged** | **0.452 ± 0.068** | **6.622** | **<0.001** |
| **Oak – unmanaged** | **0.431 ± 0.137** | **3.143** | **0.017** |
| *Tree species composition* | | | |
| Spruce – Pine | 0.065 ± 0.047 | 1.368 | 0.650 |
| **Spruce – Beech** | **0.906 ± 0.031** | **29.250** | **<0.001** |
| **Spruce – Oak** | **0.942 ± 0.058** | **16.360** | **<0.001** |
| **Spruce – unmanaged** | **0.921 ± 0.037** | **24.601** | **<0.001** |
| **Pine – Beech** | **0.842 ± 0.036** | **23.593** | **<0.001** |
| **Pine – Oak** | **0.878 ± 0.048** | **18.403** | **<0.001** |
| **Pine – unmanaged** | **0.856 ± 0.039** | **22.083** | **<0.001** |
| Beech – Oak | 0.036 ± 0.049 | 0.736 | 0.948 |
| Beech – unmanaged | 0.014 ± 0.025 | 0.570 | 0.979 |
| Oak – unmanaged | -0.021 ± 0.051 | -0.419 | 0.994 |
| *Deadwood availability* | | | |
| Spruce – Pine | -0.064 ± 0.072 | -0.885 | 0.901 |
| Spruce – Beech | 0.007 ± 0.048 | 0.152 | 1.000 |
| Spruce – Oak | -0.152 ± 0.088 | -1.741 | 0.416 |
| Spruce – unmanaged | 0.083 ± 0.058 | 1.419 | 0.616 |
| Pine – Beech | 0.071 ± 0.054 | 1.326 | 0.677 |
| **Pine – Oak** | **-0.216 ± 0.072** | **-3.025** | **0.024** |
| Pine – unmanaged | 0.147 ± 0.058 | 2.512 | 0.100 |
| Beech – Oak | 0.145 ± 0.073 | 1.985 | 0.282 |

| | | | |
|---|---|---|---|
| Beech – unmanaged | 0.076 ± 0.038 | 1.984 | 0.279 |
| Oak – unmanaged | -0.070 ± 0.077 | -0.908 | 0.894 |
| *Tree removal* | | | |
| Spruce – Pine | 0.114 ± 0.056 | 2.054 | 0.284 |
| Spruce – Beech | 0.050 ± 0.038 | 1.311 | 0.685 |
| Spruce – Oak | -0.099 ± 0.068 | -1.454 | 0.597 |
| **Spruce – unmanaged** | **0.201 ± 0.047** | **4.295** | **<0.001** |
| Pine – Beech | -0.064 ± 0.041 | 1.549 | 0.549 |
| **Pine – Oak** | **-0.213 ± 0.056** | **-3.799** | **0.002** |
| **Pine – unmanaged** | **0.087 ± 0.045** | **1.908** | **0.332** |
| Beech – Oak | -0.149 ± 0.057 | -2.633 | 0.077 |
| **Beech – unmanaged** | **0.150 ± 0.030** | **5.009** | **<0.001** |
| **Oak – unmanaged** | **0.299 ± 0.060** | **5.009** | **<0.001** |
| *Stand maturity* | | | |
| **Spruce – Pine** | **-0.261 ± 0.061** | **-4.298** | **<0.001** |
| Spruce – Beech | -0.005 ± 0.041 | -0.118 | 1.000 |
| Spruce – Oak | -0.041 ± 0.075 | -0.549 | 0.982 |
| **Spruce – unmanaged** | **0.204 ± 0.049** | **4.163** | **0.001** |
| **Pine – Beech** | **0.256 ± 0.046** | **5.580** | **<0.001** |
| Pine – Oak | 0.220 ± 0.063 | 3.469 | 0.006 |
| **Pine – unmanaged** | **0.465 ± 0.050** | **9.240** | **<0.001** |
| Beech – Oak | -0.036 ± 0.064 | -0.572 | 0.979 |
| **Beech – unmanaged** | **0.209 ± 0.034** | **6.240** | **<0.001** |
| **Oak – unmanaged** | **0.245 ± 0.067** | **3.671** | **0.003** |

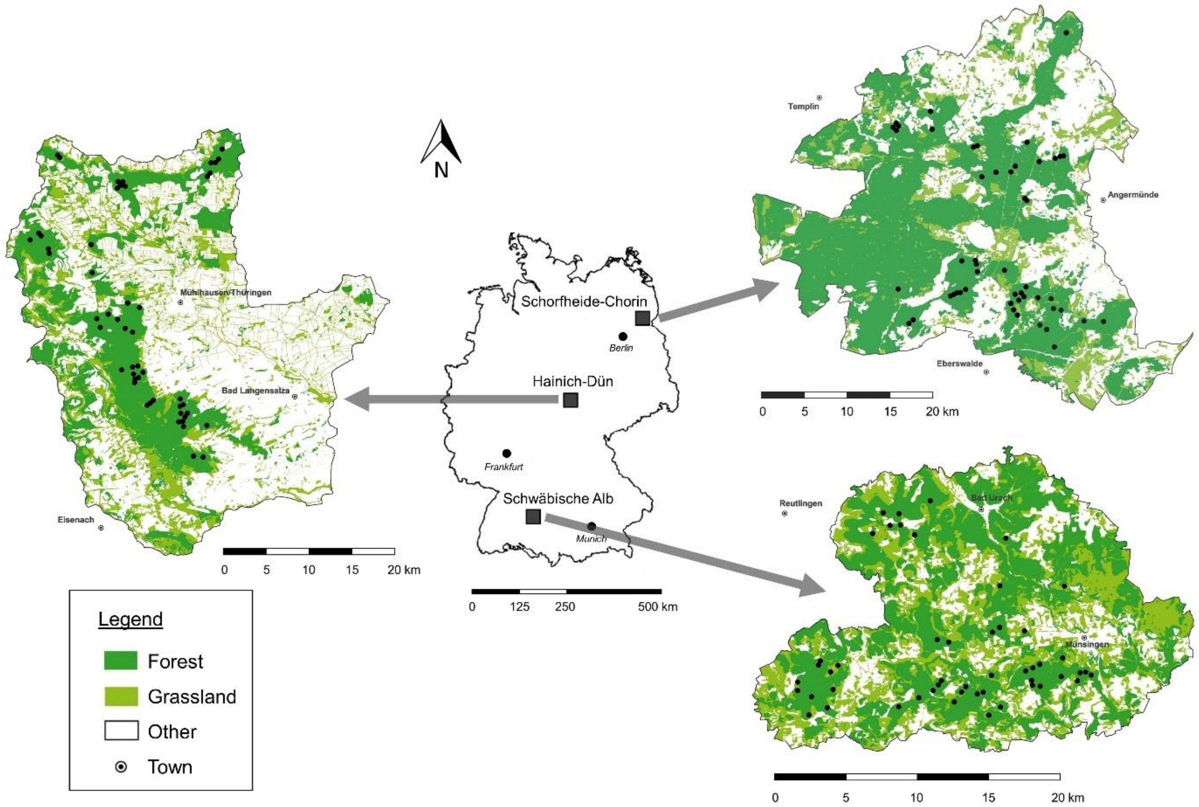

**Fig. S1** Overview of sites (indicated by black points) on which forest inventories were conducted. In three regions of Germany (Hainich-Dün: left; Schorfheide-Chorin: top right; Schwäbische Alb: bottom right) 50 plots each were inventoried. Other land use comprises agriculture, settlements and water bodies. Maps use UTM projection and are based on OpenStreetMap© data

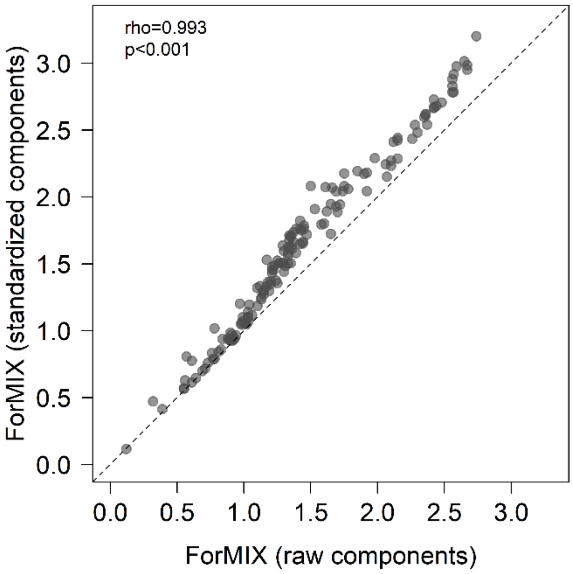

**Fig. S2** ForMIX based on raw components is highly related (Spearman's rho=0.993, p<0.001) to ForMIX for which all four components (tree species composition, deadwood availability, tree removal, stand maturity) are standardized between 0 (minimum in raw component) and 1 (maximum in raw component) before the compound index is calculated. Dashed line is the 1:1 diagonal. Data are based on forest inventories conducted on a total of 150 forest sites (size 1 ha) distributed over three regions of Germany (Fig. S1)

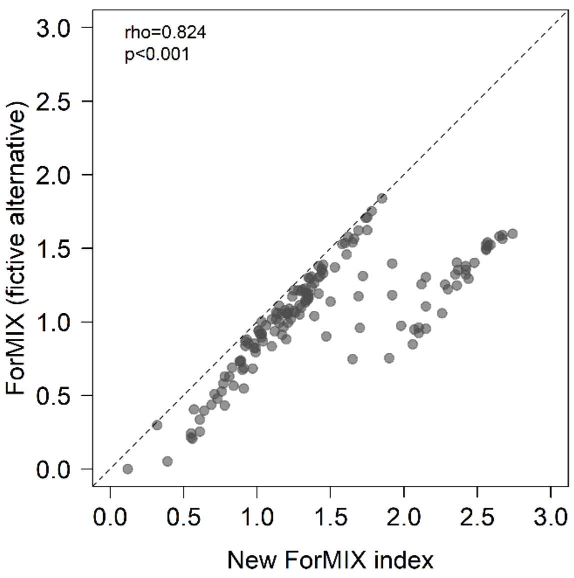

**Fig. S3** The fictive alternative ForMIX (assuming that spruce, larch and pine that jointly account for 99.5% of conifer basal area are native at all sites; expected amount of deadwood 60 $m^3$ / ha) is highly related (Spearman's rho=0.824, p<0.001) to the regular ForMIX using beech forest as reference (larch and pine not considered native, spruce only when occurring at higher elevation and accounting for less than 1/3 of basal area; expected amount of deadwood 100 $m^3$ / ha). Dashed line is the 1:1 diagonal. Data are based on forest inventories conducted on a total of 150 forest sites (size 1 ha) distributed over three regions of Germany (Fig. S1)

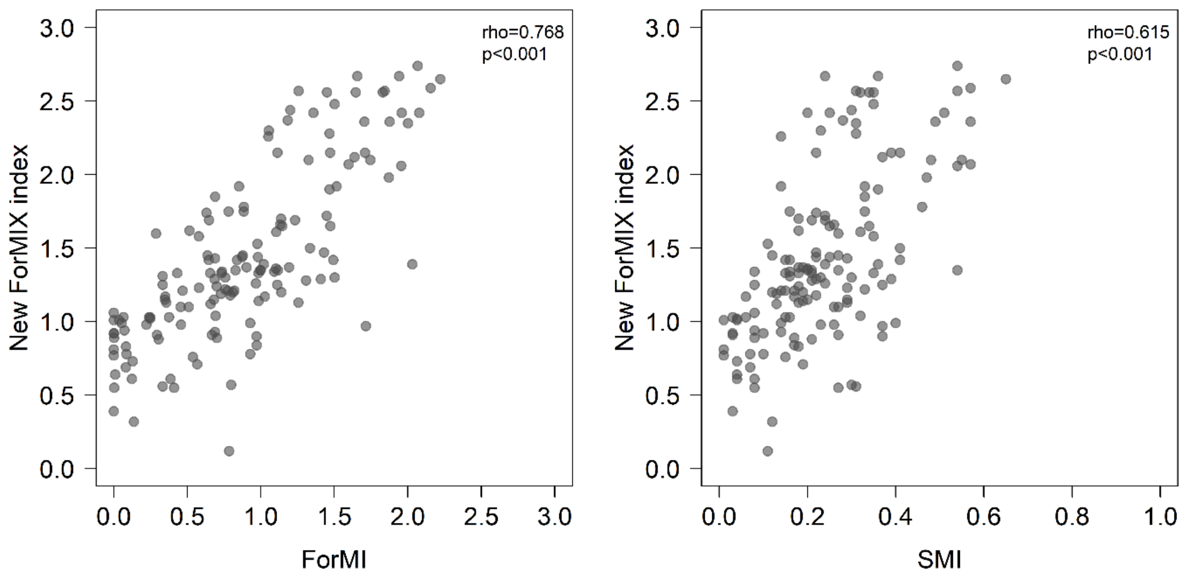

**Fig. S4** The ForMIX index correlates with alternative measures of land use in forest. Shown are relationships with the Forest Management Intensity index (ForMI, Kahl & Bauhus 2014, left) and the Silvicultural Management Intensity indicator (SMI, Schall & Ammer 2013, right), which are both highly significant (based on Spearman's rho). Data are based on forest inventories conducted on a total of 150 forest sites (size 1 ha) distributed over three regions of Germany (Fig. S1)
